# Evolution of the HIV-1 RRE during natural infection reveals key nucleotide changes that correlate with altered structure and increased activity over time

**DOI:** 10.1101/483511

**Authors:** Chringma Sherpa, Patrick E. H. Jackson, Laurie R. Gray, Kathryn Anastos, Stuart F. J. Le Grice, Marie-Louise Hammarskjold, David Rekosh

**Author notes:** Address correspondence to: David Rekosh, or Stuart Le Grice,. C.S. and P.E.H.J. contributed equally to this work.

## Abstract

**Abstract:** The HIV-1 Rev Response Element (RRE) is a *cis*-acting RNA element characterized by multiple stem-loops. Binding and multimerization of the HIV Rev protein on the RRE promotes nucleocytoplasmic export of incompletely spliced mRNAs, an essential step in HIV replication. Most of our understanding of the Rev-RRE regulatory axis comes from studies of lab-adapted HIV clones. However, in human infection, HIV evolves rapidly and mechanistic studies of naturally occurring Rev and RRE sequences are essential to understanding this system. We previously described the functional activity of two RREs found in circulating viruses in a patient followed during the course of HIV infection. The “early” RRE was less functionally active than the “late” RRE despite differing in sequence by only four nucleotides. In this study, we describe the sequence, function, and structural evolution of circulating RREs in this patient using plasma samples collected over six years of untreated infection. RRE sequence diversity varied over the course of infection with evidence of selection pressure that led to sequence convergence as disease progressed. An increase in RRE functional activity was observed over time, and a key mutation was identified that correlates with a major conformational change in the RRE and increased functional activity. Additional mutations were found that may have contributed to increased activity as a result of greater Shannon entropy in RRE stem-loop II, which is key to primary Rev binding.

**Importance:** HIV-1 replication requires interaction of the viral Rev protein with a *cis-*acting regulatory RNA, the Rev Response Element (RRE), whose sequence changes over time during infection within a single host. In this study, we show that the RRE is subject to selection pressure and that RREs from later time points in infection tend to have higher functional activity. Differences in RRE functional activity are attributable to specific changes in RNA structure. Our results suggest that RRE evolution during infection may be important for HIV pathogenesis and that efforts to develop therapies acting on this viral pathway should take this into account.

## INTRODUCTION

All retroviruses produce mRNAs that retain introns, and these mRNAs must be exported to the cytoplasm for packaging into viral particles and translation of essential viral proteins. Eukaryotic cells have RNA surveillance mechanisms that would normally restrict nucleocytoplasmic export and translation of these mRNA species (1). Thus, retroviruses have evolved specific mechanisms to overcome this restriction (2). In HIV, the Rev protein, in conjunction with its RNA binding partner, the Rev Response Element (RRE), mediates this important function (3-6).

HIV Rev contains several well-characterized functional domains that facilitate shuttling between the nucleus/nucleolus and the cytoplasm by accessing cellular pathways for nuclear import and export. One of these domains is a basic, arginine-rich motif that functions as a nuclear/nucleolar localization signal (7-10). This domain also serves as an RNA-binding domain that binds specifically to the RRE (11, 12). It is flanked on both sides by oligomerization domains that are required for Rev multimerization on the RRE (13, 14). Towards the carboxy terminus is a leucine-rich domain which docks on the Crm1-RanGTP complex (15) and functions as the nuclear export signal (NES) (12, 16, 17).

The RRE is a *cis*-acting RNA element located in a highly conserved and functionally important region of *env* at the N-terminus of the gp41 fusion protein (3-6). At this position, it is present in all HIV RNAs that retain introns. The minimal functional RRE, or “short” RRE, has been mapped in infectious laboratory clones to a 234-nt region (see Figure 3). It forms a highly branched structure where the 5’ and 3’ ends pair to form stem-loop I (SL-I) (18). SL-I opens into a central loop, from which several additional stem loops branch out. Stem loop II contains stem IIA that branches out of the central loop and opens into a three-way junction. The junction opens into stem-loops IIB and IIC (19, 20). SL-III is a small stem-loop that comes off of the central loop, while SL-IV and SL-V can adopt alternative topologies (21). Maximal functional activity of the RRE requires a somewhat larger structure, often referred to as the “long” (351-nt) RRE, characterized by an extended SL-I (18, 22, 23).

Primary RRE binding initiates cooperative assembly of additional Rev molecules (about 6-8) in a process that requires electrostatic Rev-RRE interactions and hydrophobic Rev-Rev interactions (13, 18, 24-32). Rev oligomerization on the RRE increases its binding affinity ~500 fold (30). Oligomerization also arranges the NES domains for binding to the RAN-GTP bound Crm1 dimer (15) forming an export-competent ribonucleoprotein complex (33, 34). The complex is targeted to the nuclear pore, where it interacts with the nucleoporins, resulting in its translocation to the cytoplasmic side. Once in the cytoplasm, RRE-containing mRNAs are translated into Gag, Gag-Pol, Env, and accessory viral proteins.

Molecular details of the Rev-RRE pathway have been delineated mostly from studies of lab-adapted HIV clones. However, there is mounting evidence that subtle variation in Rev and RRE sequences among primary isolates may contribute to pathogenesis. Differences in Rev-RRE functional activity up to 24-fold were observed for naturally occurring viruses in different patients (35). In addition, a study performed with a Thai cohort demonstrated RRE changes over the course of infection, and higher RRE activity was associated with a more rapid CD4 count decline (36). Low Rev activity has also been associated with slower disease progression (37, 38) and reduced susceptibility of infected cells to T-cell killing (39). In equine infectious anemia virus, a related lentivirus that also utilizes the Rev/RRE axis, functional evolution of Rev has been observed during infection, and Rev activity has been found to correlate with disease state in ponies (40-42). Further studies of the structural and functional evolution of the Rev-RRE system in natural infection are necessary to understand the role that this regulatory axis plays in adaptation of HIV to diverse immune environments, and this may benefit development of Rev-RRE targeted therapeutics.

We previously investigated the activity of Rev-RRE cognate pairs from HIV isolated from five different patients at two time-points during their course of infection (43). The sequences were obtained by single genome sequencing of viruses from blood plasma samples and their functional activity was determined using a sub-genomic reporter assay. We observed significant activity differences between Rev-RRE cognate pairs from different patients and from different time points in the same patient. Evolution of Rev and the RRE observed in patient SC3 was particularly striking. In this patient, the RRE converged on one predominant sequence at the later time point (the M57A RRE, designated here as V20-1), suggesting it was subject to strong selective pressures. Furthermore, this RRE had only four nucleotide changes (mut 1-4) relative to the early time point RRE (M0-A RRE, designated here as V10-2) but was 2-3 fold more functionally active. Gel mobility shift assays revealed that the V20-1 RRE promoted Rev multimerization at a lower concentration of Rev protein compared to the V10-2 RRE. It was also notable that the predominant Rev sequence present at the early time point persisted at the late time point, suggesting that the limited nucleotide changes in the RRE were the major driver of differential Rev-RRE activity.

In the current study, we conducted a detailed examination of RRE evolution in the blood plasma of patient SC3 using samples collected at six-month intervals, from the time viral RNA was first detected through year six of infection. DNA deep sequencing, selective 2′ hydroxyl acylation analyzed by primer extension (SHAPE) chemical probing, and a Rev-RRE functional assay were used to determine sequence evolution, RRE secondary structure, and Rev-RRE functional activity over time and to explore the mechanism underlying the observed activity differences. This study highlights, for the first time, the structure-function relationship of longitudinal RRE sequence evolution in a single patient and the underlying molecular mechanisms.

## RESULTS

### RRE sequence evolution in an HIV-infected individual followed over many years

Plasma samples from patient SC3 were collected over a period of 10 years during standardized visits spaced about 6 months apart as part of the Women’s Interagency HIV Study (WIHS) Consortium in the Bronx, New York. Patient SC3 was enrolled in the WIHS cohort prior to HIV seroconversion, was followed through visit 20, and never received antiretroviral therapy (ART). Samples for this patient were available for all time points with the exception of four missed visits (2, 3, 17 and 18). Plasma samples from each WIHS visit were tested for p24 antibody and CD4 count (Figure 1). Viral load was measured once the patient tested positive for p24 antibodies. In this report, each visit is denominated as VXX with the numeral representing the visit number. Although p24 antibodies were not detected under the original WIHS protocol until V10, a significant fall in her CD4 count was noted at V08 and we were able to readily amplify HIV from plasma taken at this time point, suggesting infection between V07 and V08. After a short-lived rebound at V09, the CD4 count continued to fall and reached a nadir of 73 cells/μl at V20. Seroconversion and viremia were noted in the data obtained from the WIHS protocol at V10, and the viral load continued to rise through V20 reaching a peak of 1.4 × 10^6^ copies/ml. As expected without ART treatment, we observed a general increase in viral load concomitant with a decline in CD4 count as disease progressed.

**Figure 1:**
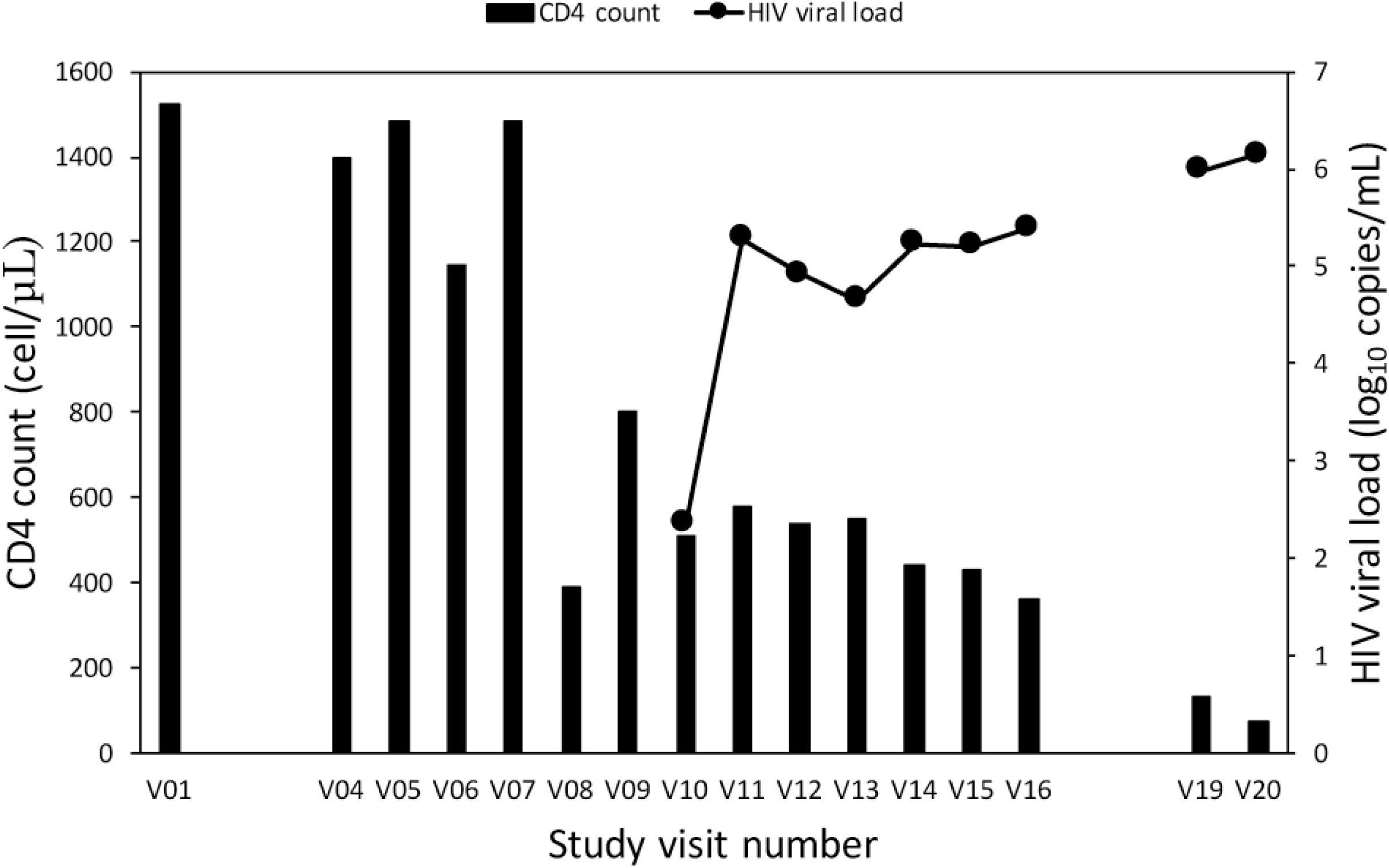
Patient SC3 HIV viral load and CD4 count at each visit during participation in the WIHS cohort. Sequential study visits occurred about six months apart and blood samples were obtained at each visit. V01 refers to the first visit, V04 to the fourth, etc. Data is missing for unattended visits. HIV sequences could first be amplified from plasma samples obtained at V08 and onwards. However, HIV seroconversion was first appreciated under the WIHS protocol at V10 and viral loads were only performed from this visit on. The patient died after V20.

We investigated RRE sequence evolution in viruses isolated from V08 and onwards. A total of 12 plasma samples representing visits 7, 8, 9, 10, 11, 12, 13, 14, 15, 16, 19 and 20 were received from the WIHS consortium. Viral RNA isolation was attempted from each sample and, if successful, RRE sequences were determined by next-generation sequencing. As stated above, the first visit to yield HIV sequences that could be amplified by PCR was V08. The prevalence of each RRE sequence at a given visit was calculated as a percentage of the total number of sequences present at that time point (Figure S1). Sequences present with a frequency of <5% were then excluded from consideration, as it was not possible to distinguish true rare RREs from artifacts due to PCR errors, and the prevalence of significant variants was recalculated after exclusion of these minority sequences (Figures 2, S2). RRE sequences were assigned a code in the form VXX-Y where XX refers to the visit number and Y refers to the prevalence rank order of that sequence.

**Figure 2:**
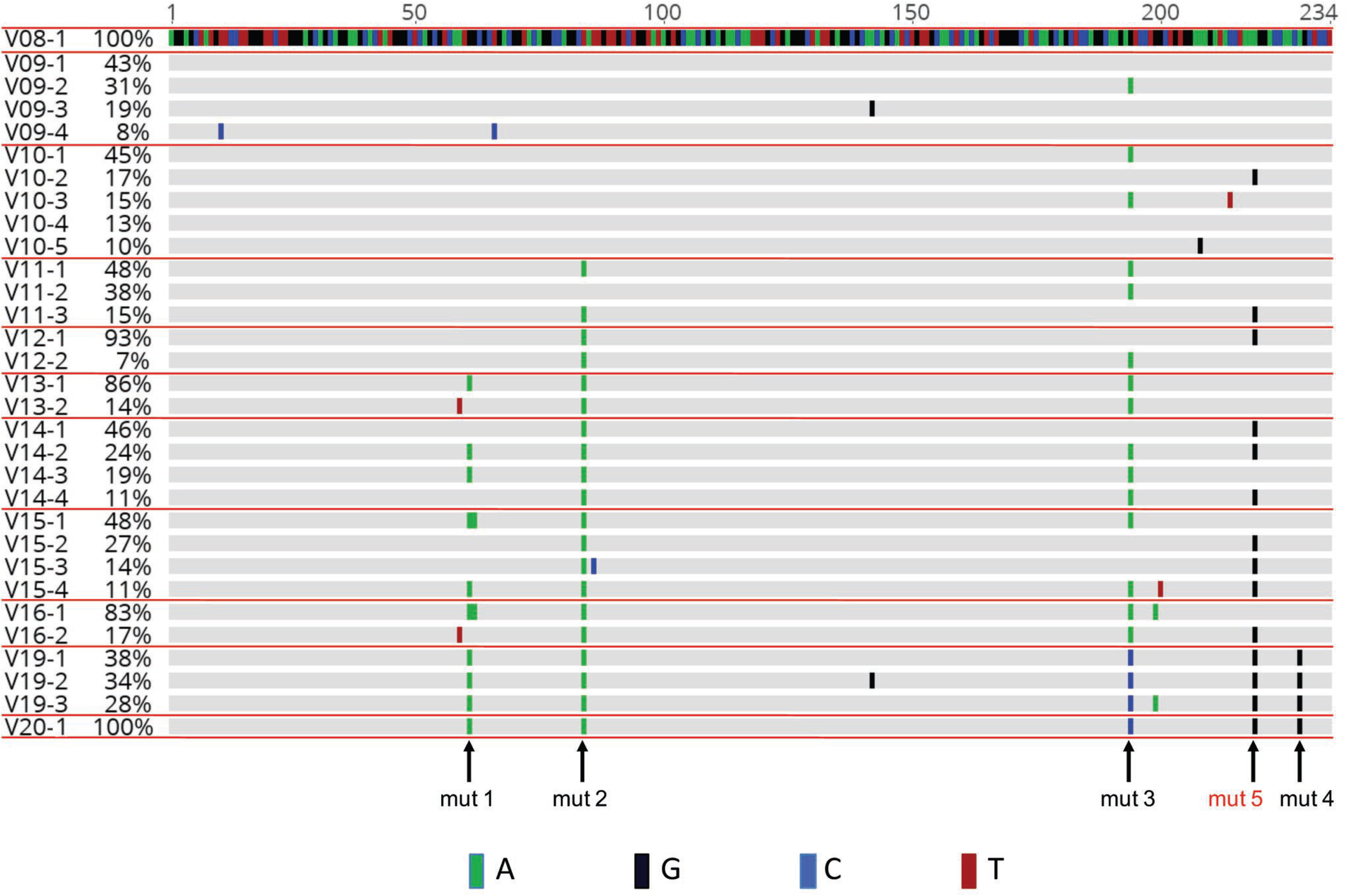
Evolutionary alignment of SC3 RREs. Deep sequencing of viral RNA was performed from plasma collected at each study visit. This alignment includes RRE contigs that were present in at least 5% of the total sequences as explained in the methods. Sequence labels are in the form of VXX-Y where XX refers to visit number and Y refers to rank order of the contig within that visit plasma sample. Nucleotide changes relative to the presumptive founder sequence, V08-1, are highlighted. Mutations (Mut) 1-5 refer to single nucleotide changes that occur in V20-1 relative to V08-1. Mut 5 is highlighted in red, as this change is also found in V10-2 and was not recognized in our previous study of the “early” and “late” RREs from patient SC3 (43). (See also Figures S1 for the alignment including minority variants and S2 for the complete nucleotide sequences included here.)

As shown in Figure 1, data obtained from the WIHS indicated that patient SC3 seroconverted to HIV-positive status at V10. Thus, in our previous study, V10 was believed to represent a timepoint within six months of infection and was designed month 0 (M0). An RRE was identified at this visit by single genome sequencing that was believed to be the major species present at that time and designated an “early” RRE. Our deep DNA sequencing results showed that this “early” RRE sequence represented 17% of the sequences present at that visit and it is therefore labeled as V10-2 in the present study. This sequence is very similar to the single founder RRE sequence found at V08 (V08-1), differing only by a single nucleotide near the 3’ end and mapping to the right-hand side of SL-1 (designated here as mut 5). Our previous study also identified a single RRE from V20 which we designated a “late” RRE. This RRE is identical with the V20-1 sequence generated by deep sequencing.

Since the V10-2 RRE previously described as the “early” RRE has been studied intensively by us, we decided to compare its properties to the founder V08-1 RRE. Both RREs were transcribed *in vitro* and their secondary structures chemically probed by 1M7 (1-methyl-7-nitroisatoic anhydride) using selective 2′ hydroxyl acylation analyzed by primer extension and mutational profiling (SHAPE-MaP) (44-50). In this procedure, RNAs are first modified with chemical reagents that selectively acylate unpaired ribonucleotides at their 2′-hydroxyl positions, then reverse transcribed under conditions that introduce mutations in cDNA opposite the sites of modification (47, 49). The position and frequencies of these mutations are used to create reactivity profiles indicating which RNA nucleotides are likely to be single- or double-stranded (data available upon request). This information is in turn used to guide the RNAstructure software (51) to generate lowest Gibbs free energy secondary structure models.

SHAPE-MaP studies revealed that V08-1 and V10-2 form similar structures with only minor local differences (Figure 3A and 3B). These structures are similar to the previously reported ARv2/SF2 RRE structure (22) where a part of the central loop (nt 125-130) between SL-III and SL-IV paired with nucleotides from the upper stem of SL-I (193-198) forming a stem that bridges the central loop and SL-IV and SL-V.

**Figure 3:**
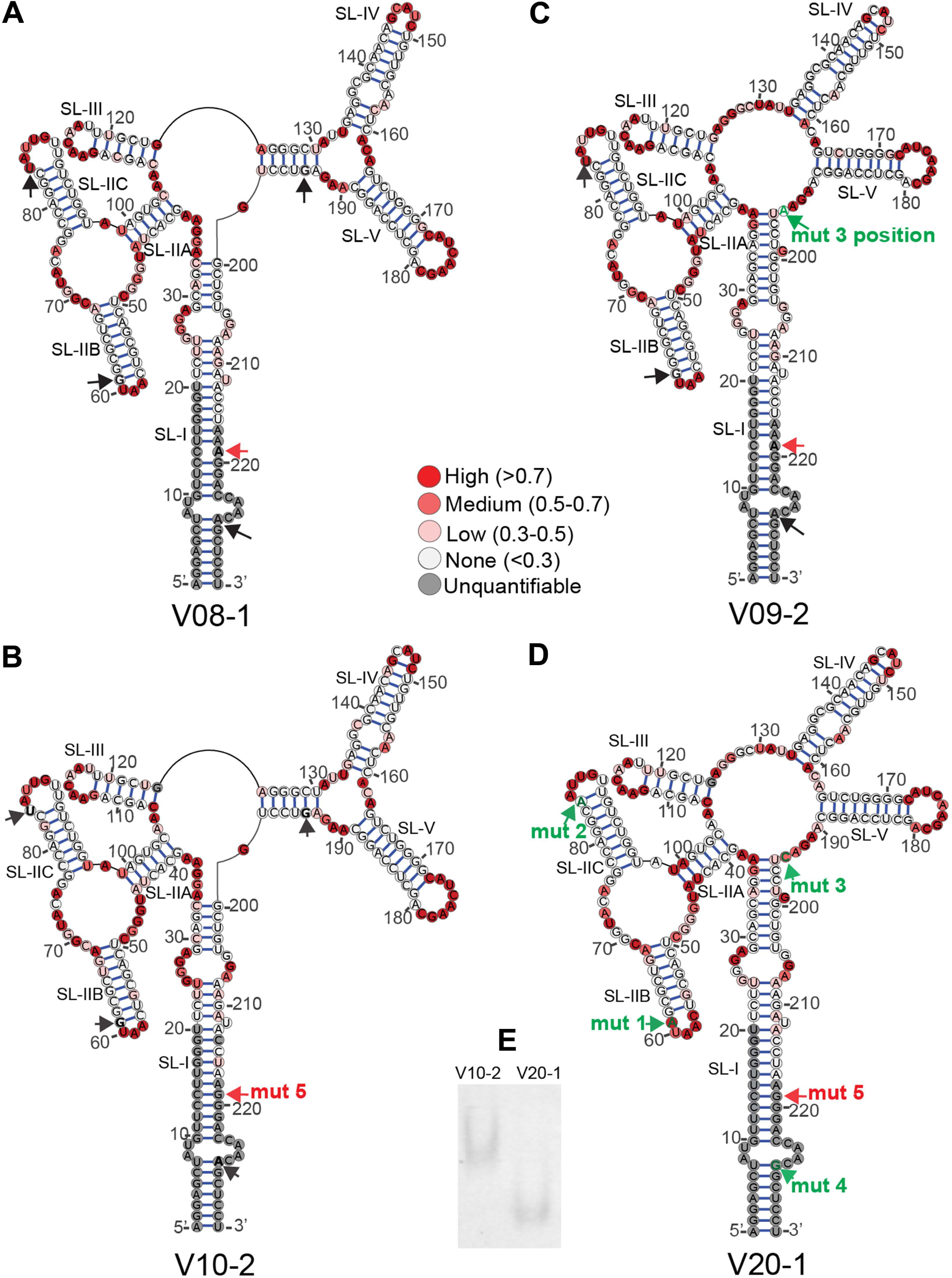
Structures of selected SC3 RREs. Secondary structures of [A] V08-1, [B] V10-2, [C] V09-2, and [D] V20-1 234-nt RREs were determined by SHAPE-MaP. 1M7 reactivities of RRE nucleotides are color coded and superimposed on the structures. The positions of the mut 1, mut 2, mut 3, and mut 4 single nucleotide changes are represented by black arrows. Green labelled arrows show where the nucleotide at this position varies from V08-1, the presumptive founder sequence, and V10-2. In V09-2, the mut 3 position shows a G to A change, while the mut 3 position in V20-1 shows a G to C change. The mut 5 position is shown with a red arrow and is additionally labelled with “mut 5” in structures where the nucleotide varies from V08-1. [E] Migration of V10-2 and V20-1 RRE on a native PAGE gel was visualized after 22 hours by UV shadowing.

To determine activity of each of these RREs, as well as V20-1, each RNA was cloned into a two-color fluorescent-based proviral reporter vector (52) which carried inactivating mutations in *rev*. RRE activity was measured by co-transfecting a separate plasmid that expressed the previously described M0-B/M57A-SC3 Rev into 293T/17 cells. This Rev protein has been shown by single genome sequencing to pair with the V10-2 RRE (43). It was also the unique founder Rev sequence present at V08. Unexpectedly, this sequence still persisted at V20 and was found in single genomes with the V20-1 RRE (43). The reporter construct expressed eGFP from the intron-containing Rev-dependent *gag* mRNA and TagBFP from the intron-less Rev-independent *nef*-like transcript. Thus, in this assay, the ratio of the fluorescent signals (eGFP:TagBFP) is a measure of Rev-RRE functional activity. Control experiments showed that the assay is highly sensitive to the addition of Rev. For example, when increasing the amount of Rev plasmid in the assay system from 0 ng to 100 ng, there is a 268-fold increase in GFP expression with only a slight decrease in BFP expression (2.6 fold) (52).

All three RREs displayed clear Rev responsiveness as the fluorescent ratio in the non-Rev containing transfections was at least 500-fold less than in Rev-containing transfections (data not shown). Despite their local structural differences, both V08-1 and V10-2 RRE showed similar levels of RRE activity (*p*=0.999) (Figure 4). Therefore, although mutation 5 arose after infection and increased in prevalence through V20, it did not increase RRE functional activity or significantly change RRE structure. Additionally, activities of V08-1 and V10-2 RREs were substantially less than that of V20-1 (Figure 4) (*p*=0.022 and *p*=0.021, respectively), consistent with our previous finding that V20-1 RRE is more active (43). Notably, we also replicated the finding that V20-1 is significantly more active than V10-2 using a lentiviral vector system, where packaging of genomic RNA and titer is dependent on Rev-RRE function (data available upon request) (35, 52).

**Figure 4:**
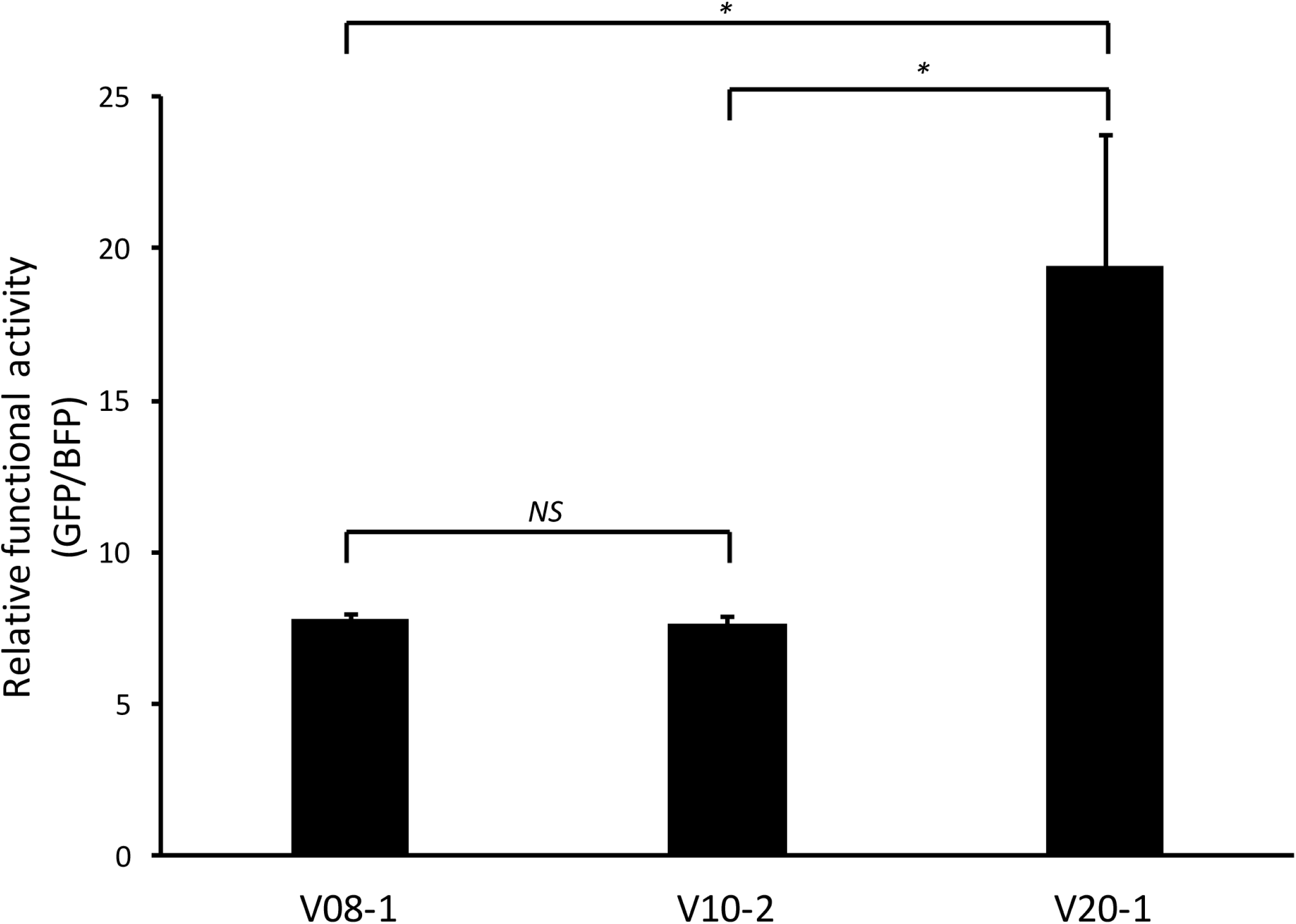
Functional activity of SC3 founder (V08-1), “early” (V10-2), and “late” (V20-1) RREs. Functional activity of the three RREs was determined by transfecting reporter constructs containing the different RREs into 293T/17 cells along with 50 ng of SC3 Rev and measuring the ratio of the mean fluorescent intensity of GFP to BFP. *N*=4, SEM is represented by error bars. P-values of <0.05 are represented by *, NS=non-significant difference.

### V10-2 and V20-1 RREs form distinctly different secondary structures

We previously reported that there are only four single nucleotide changes (mut 1-4) between the V10-2 and V20-1 RREs (43). A fifth mutation (mut 5) is present in both V10-2 and V20-1 and therefore did not score as a difference between the two RREs identified as “early” and “late” in our previous study. On a canonical 234-nt five stem-loop (5SL) RRE structure (21), these changes map near the apical loop of SL-IIB [nt 61: mutation (mut) 1] and SL-IIC [nt 84: mutation (mut) 2], the central loop region between SL-V and SL-I [nt 194: mutation (mut) 3], and at the base of stem of SL-I [nt 228: mutation (mut) 4] (see Figures 3B and 3D). We next explored whether the differences in activity between the two RREs could be attributed to structural differences caused by any individual nucleotide change.

*In vitro* transcribed and folded V10-2 and V20-1 short RREs migrated at a significantly different rate when analyzed by native polyacrylamide gel electrophoresis (Figure 3E), suggesting major structural differences. Consistent with the gel migration data, SHAPE-MaP of the two 234-nt RREs revealed that the V10-2 and V20-1 RREs adopt distinctly different conformations (Figures 3B and 3D). Unlike V10-2, which had part of the central loop region base-paired, V20-1 formed the canonical 5SL structure where five distinct stem-loop structures (SL-I to SL-V) radiate out directly from the single stranded central loop.

Closer analysis of the location of each of the four nucleotide changes suggested that the major structural shift in the two RREs might have been caused by mut 3. The nucleotide at the mut 3 position (G) is base-paired in the bridging stem in V10-2. In V20-1, it mutates to C disrupting base pairing at this position thereby destabilizing the stem formed by nt 125-130 and 193-198. Consequently, nt 34-36 were significantly more reactive while nt 125, 127, and 130 were significantly less reactive in V10-2 compared to V20-1. Mut 1 and mut 2 produced more limited local structural changes, i.e. opening and formation of the base-pair near the apical loop of SL-IIB and SL-IIC in V20-1 relative to V10-2. Since mut 4 was located at the primer-hybridization site (see the PCR1 step of SHAPE-MaP in Methods) for the 234-nt RREs, we were unable to assess its effect on the secondary structure.

A 3-dimensional structural analysis of a 351-nt RRE by small angle X-ray scattering has shown that the extended SL-I folds back on regions in and around SL-II to expose a cryptic Rev-binding site (22). Data obtained in connection with our previous study (43), showed that the extended SL-I of the 351-nt V10-2 and V20-1 RREs has an additional four nucleotide changes at nucleotide positions 13, 27, 29, and 57 (using the 351-nt RRE numbering system). We therefore investigated if these nucleotide changes in the extended SL-I affect RRE structure. Additionally, probing these extended RREs provided an opportunity to determine the effect of mut 4 on their structure, as this position no longer fell on the primer hybridization site during SHAPE analysis of the 351-nt RREs. The long RREs were investigated by capillary electrophoresis-based SHAPE (Figure 5) (CE-SHAPE) using N-methylisatoic anhydride (NMIA) as the electrophile (54, 55). CE-SHAPE differs from SHAPE-MaP in that the chemically modified nucleotides are identified as stops during primer extension. CE-SHAPE analysis of the 351-nt V10-2 and V20-1 revealed that mut 4 did not produce any structural changes between V10-2 and V20-1 RREs, as it changed a U-A base pair to U-G. Additionally, the nucleotide differences in the extended SL-I region of these RREs produced only highly localized structural changes, if any. Changes at nt 13 and nt 57 produced no structural changes at all, whereas nucleotide differences at nt 27 and nt 29 induced opening of base pairs at these positions in V20-1. As these mutations in SL-1 did not alter the structure of the comparable region of the 234-nt RRE, subsequent studies investigated their effects only in the short RRE forms.

**Figure 5:**
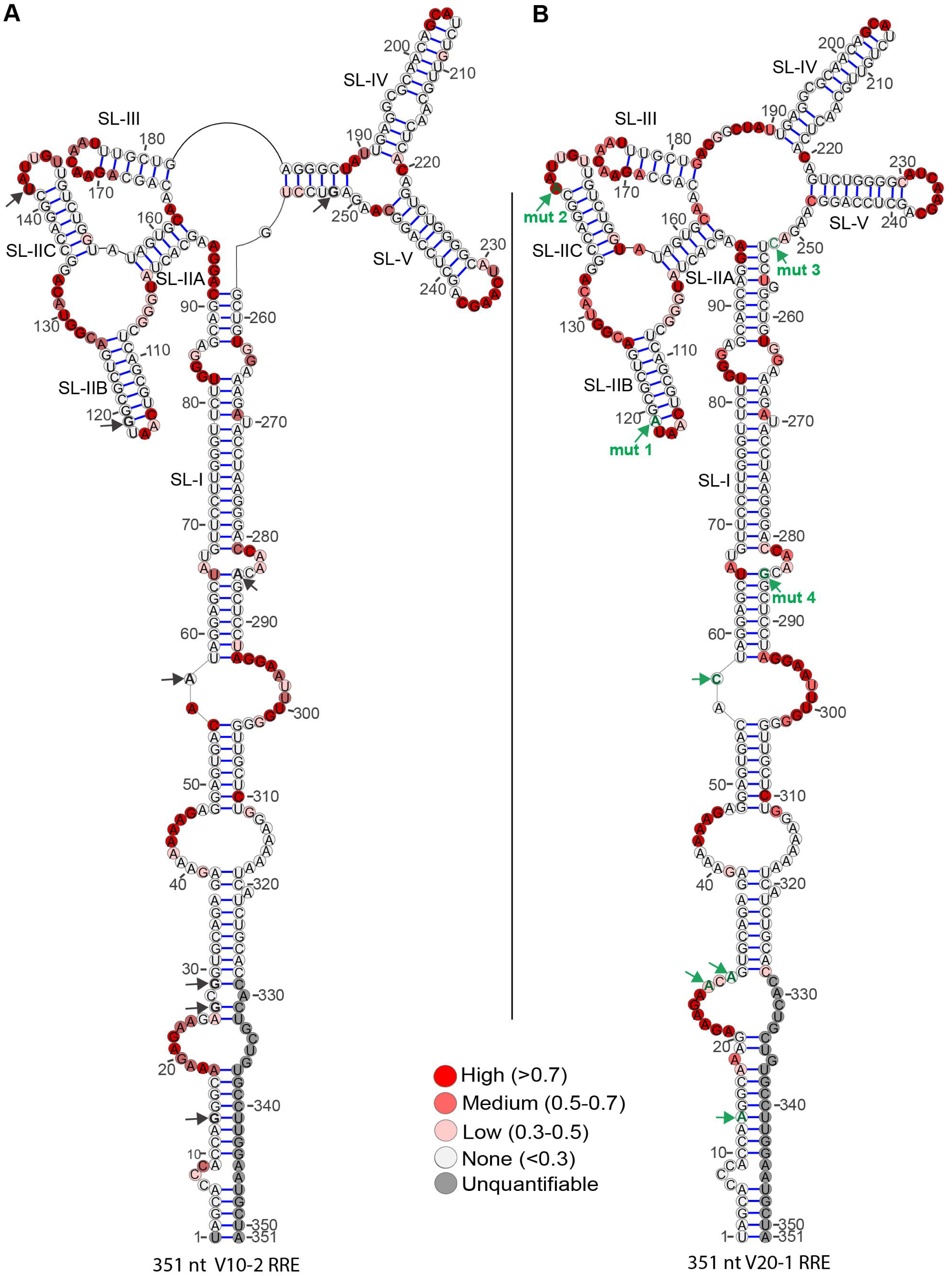
Secondary structures of full-length (351-nt) V10-2 and V20-1 SC3 RREs. Secondary structures of [A] V10-2 and [B] V20-1 SC3 RREs, determined by CE-SHAPE. NMIA nucleotide reactivities are color coded and superimposed on the structures. The positions of the eight nucleotides differing between the two RREs are represented by arrows. Mutations within the short 234-nt RRE are designated mut 1, mut 2, mut 3, and mut 4 and shown in green on the V20-1 RRE. Four additional mutations within stem SL-I are shown as green arrows without labels on the V20-1 RRE. The position of each of these changes is shown by black arrows on the V10-2 RRE.

### Structural and functional analysis of patient SC3 RREs containing individual single nucleotide mutations

We next investigated the contribution of each of the four nucleotides which differed between the V10-2 and the V20-1 RREs to alterations in structure and functional activity. To this end, we created four new synthetic 234-nt RRE sequences, each containing one of the single nucleotide changes in a V10-2 RRE background. The new RREs containing mut 1, 2, 3, and 4 were respectively termed M1, M2, M3, and M4. The secondary structure of each RRE was probed by SHAPE-MaP (Figure 6A-D). As predicted, M1, M2, and M4 RREs formed V10-2 like structures, whereas M3 RRE formed a V20-1 like structure. M1 RRE resembled V10-2 RRE with the exception that the G to A mutation eliminated a base pair adjacent to the SL-II apical loop. Similarly, M2 RRE differed from V10-2 RRE only in the formation of an extra base pair next to the apical loop of SL-IIC. M4 RRE formed a structure identical to V10-2 RRE, since the mut 4 A to G change preserved non-canonical base pairing at this position by replacing A-U with a G-U base pair. The major structural shift involving the central loop, SL-IV, SL-V, and SL-I between the V10-2 and V20-1 RREs was reproduced by the single G to C mutation in the M3 RRE. Therefore, we reasoned that mut 3 might be the major driver of the functional difference between the V10-2 and V20-1 RREs.

**Figure 6:**
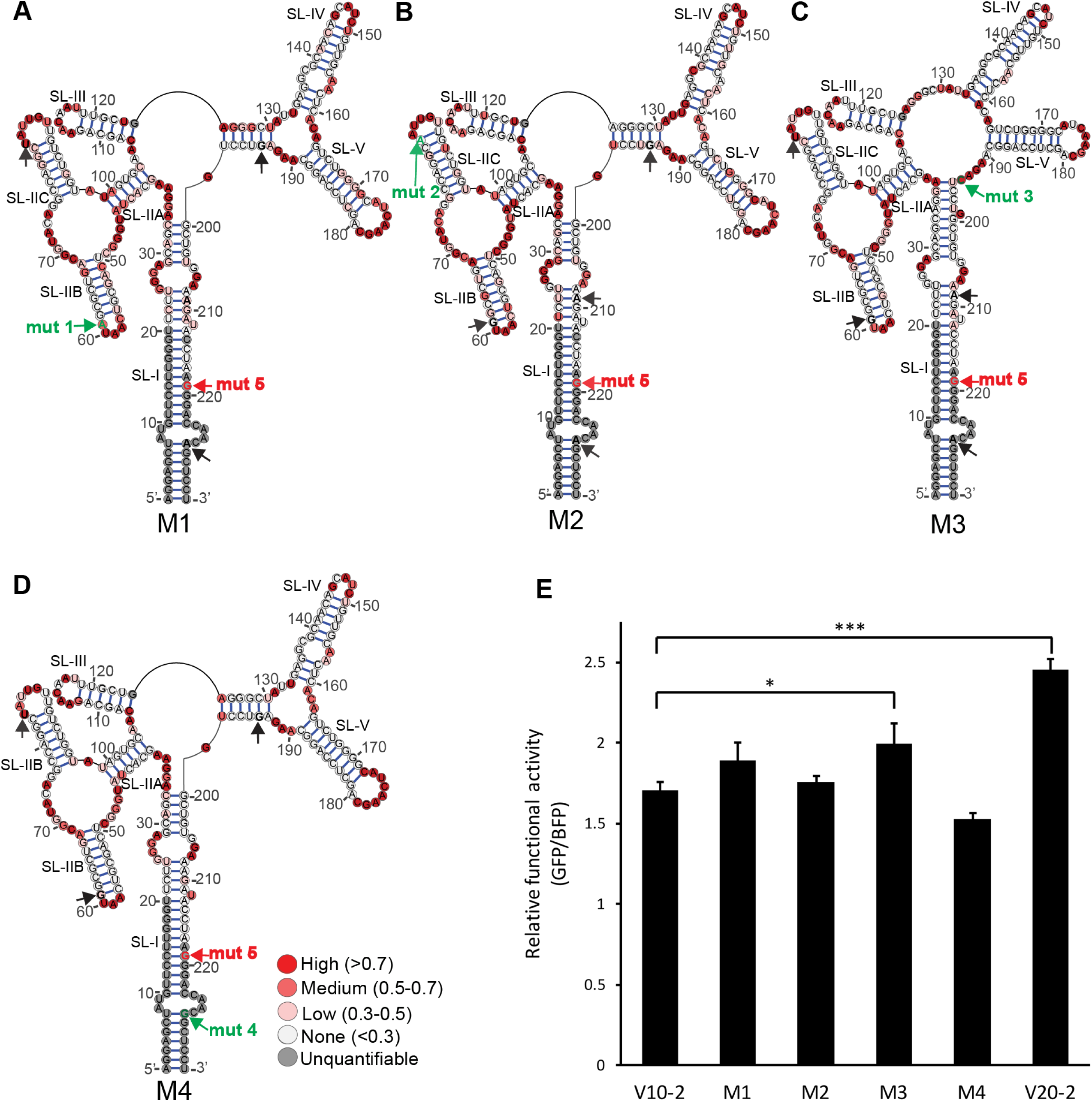
Secondary structures and function of V10-2 RRE single mutants. Secondary structures of the [A] M1, [B] M2, [C] M3, and [D] M4 SC3 RREs were determined by SHAPE-MaP. 1M7 reactivities are color coded and superimposed on the structures. The positions of mut 1-5 are represented by black and green arrows. Mut 1, mut 2, mut 3, and mut 4 are indicated in green on the RREs where the nucleotide at that position differs from V10-2, while mut 5 is represented in red and is present on every structure. [E] Rev-RRE functional activity of M1-M4 RREs compared with V10-2 and V20-1 in the presence of 100 ng SC3 Rev. N=2, SEM are represented by error bars. Selected p-values of <0.05 and <0.001 are represented by * and ***, respectively.

We next measured the contribution of each mutation to the functional activity difference between the V10-2 and V20-1 RREs in 293T/17 cells (Figure 6E). Activity of RREs M1-M4 was determined using the fluorescent-based transient proviral reporter with the same cognate SC3 Rev used in Figure 4. As before, in the presence of SC3 Rev, functional activity of V20-1 RRE was significantly higher than that of V10-2 RRE (*p*=0.001). Among all single SC3 mutants, only M3 displayed activity that was statistically significantly higher than V10-2 (*p*=0.072) even though it did not quite reach the activity level of V20-1. M1 and M2 RREs were only slightly more active than V10-2 but these differences did not reach statistical significance. Activity of M4 RRE was also not statistically distinguishable from that of V10-2. However, M4 had a tendency towards lower, rather than higher, functional activity. Thus, the difference in functional activity between V10-2 and V20-1 could not be explained by any single nucleotide mutation.

### Structural and functional analysis of selected RRE sequences derived from patient SC3 at different time points

To further understand how specific sequence changes between V10-2 and V20-1 RREs affected functional activity, we studied intermediate RRE sequences from viruses that arose during disease progression in patient SC3. SC3 RREs that allowed us to test combinations of these mutations in the order they appeared in the natural isolates were selected from this evolutionary data set (Figure 2; Figure S2). The secondary structure and functional activity of these RREs were then determined by SHAPE-MaP and the fluorescent-based reporter assay, respectively.

The position designated mut 3 starts as a G in V08-1, remains a G in V10-2, changes to A in many sequences in V09 through V16, and then to C in V19 and V20. As this is the position that causes the major structural shift between V10-2 and V20-1, we sought to determine if the A at the mut 3 position would result in the same structure as its replacement by C. To test this, we determined the secondary structure of V09-2 RRE by SHAPE-MaP and compared it to the structure of V08-1 and V10-2 (Figure 3A-C). The V09-2 RRE sequence is identical to V08-1 except at the mut 3 position, where the G in V08-1 is replaced with A. We hypothesized that if disruption of the G-C base-pair at this position induced the structural shift observed between V10-2 and V20-1, then such a shift should also be reflected between V08-1 and V09-2. Indeed, the secondary structure of V09-2 resembled that of V20-1 RRE, confirming our hypothesis. V09-2 RRE was also slightly more active than V08-1 RRE (though this trend did not reach statistical significance) (*p*=0.116) and V10-2 (*p*=0.027) (Figure 7). Consistent with our previous observation with M3 RRE, the structural shift invoked in V20-1 and V09-2 relative to V08-1 and V10-2 shows a trend towards increased SC3 RRE activity.

**Figure 7:**
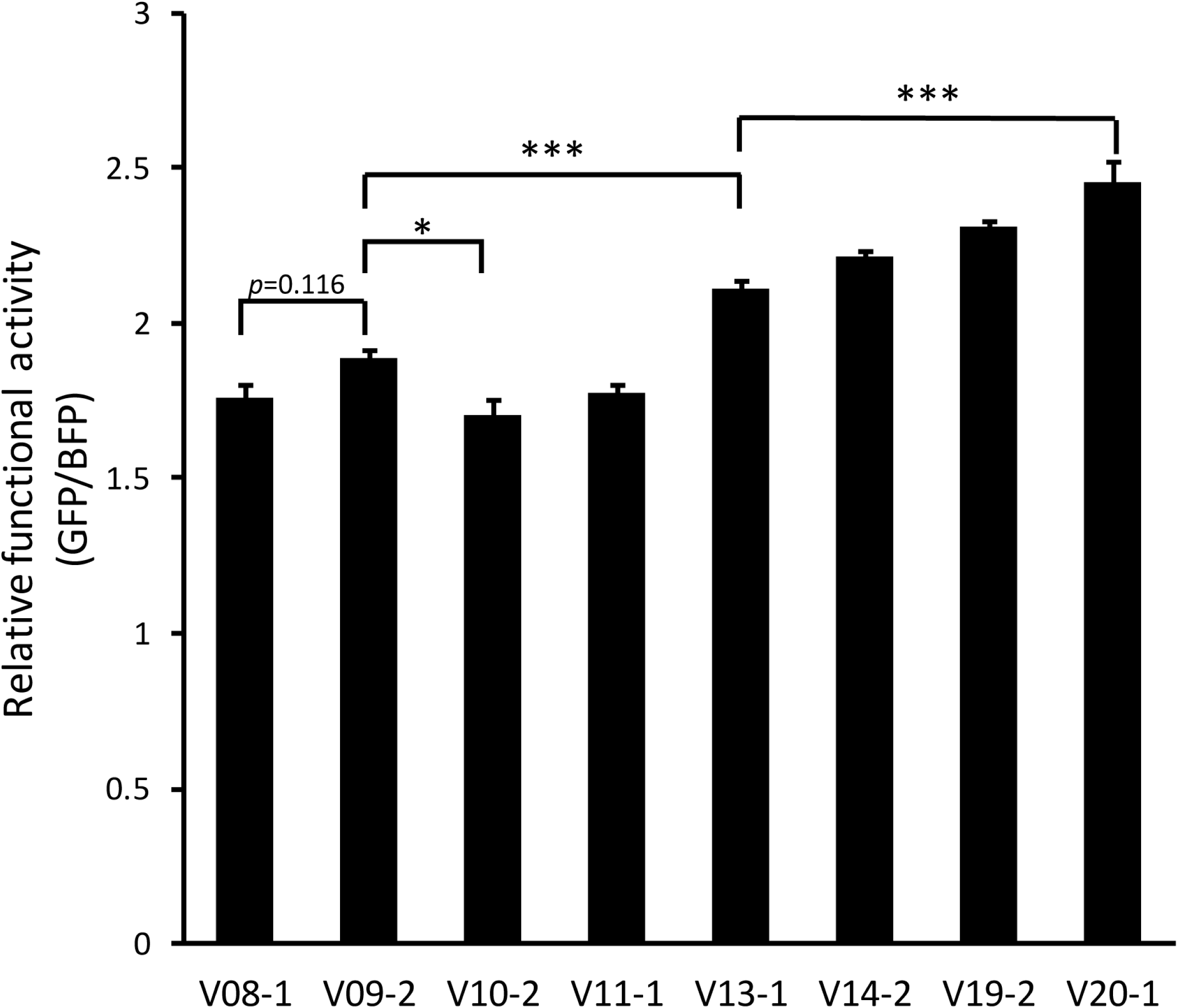
Rev-RRE activity of SC3 RRE haplotypes. Activity of selected RREs was determined in the presence of 100 ng Rev using the fluorescent assay system. N=2 except for V9-2 and V13-1 where N=4. SEM are represented by the error bars. Selected p-values of <0.05 and <0.001 are represented by * and ***, respectively.

Mut 2 was first noted in the sample from V11 and was present in all RRE sequences from subsequent visits. We tested the V11-1 RRE as it provided the opportunity of investigating the combination of mut 2 together with A in the position of mut 3. Although it was predicted to have higher activity than M3 and V09-2 RRE, V11-1 RRE had similar level of RRE activity as the founder RRE, V08-1, suggesting that while A at the position of mut 3 increases functional activity alone, it does not do so in the context of the additional mut 2 (*p*=1.000).

We first detected mut 1 in plasma samples from visit 13. This mutation was frequently observed on subsequent visits and was present in the majority of RREs from visit 15 onwards. The V13-1 RRE contained not only mut 1 but also mut 2 and A in the mut 3 position. This RRE had significantly higher activity than both the V08-1 and V09-2 variants (*p*<0.001 for both comparisons). Therefore, mut 1 not only contributes to RRE activity in itself but also rescues the activity of the combination of mut 2 and an A in the mut 3 position.

We also tested the activity of V14-2, which allowed us to test the combined phenotype of mut 1, mut 2, A in the mut 3 position, and mut 5. Activity of this RRE did not significantly differ from V13-1, suggesting that acquisition of mut 5 does not contribute to RRE activity (*p*=0.332). However, activity of the V20-1 RRE, which harbors C in the position of mut 3 rather than the A of V13-1, as well the other mutations, was significantly higher than V13-1 (*p*<0.001). This suggests that C in the mut 3 position is important for imparting higher RRE activity when all other mutations are present. It is also possible that although mut 4 does not contribute to RRE activity individually, it can contribute to activity when combined with the other additional changes.

This set of SC3 RREs permitted examination of changes in functional activity over time. Activities of V08-1, V10-2, and V11-1 were very similar, demonstrating that the combinations of mutations observed in these sequences are not sufficient in themselves to confer higher functional activity. V13-1 was the first tested RRE to arise, other than V09-2, that showed significantly higher activity than V08-1 and it did so with the combination of mut 1, mut 2, and A in the mut 3 position. The accumulation of additional changes in V14-2 (addition of mut 5), V19-2 (addition of mut 4, the A to C change at mut 3, and an additional A to G change at nt 142), and V20-1 (reversion of nt 142 to A) corresponded to a trend towards increasing functional activity at each step. This steady trend suggests functional evolution of the RRE in the later phase of the disease course of patient SC3, as new combinations of mutations arose and those conferring greater functional activity showed preferential selection.

### Shannon entropy profile of SC3 RRE variants and single mutants

Since the energetically most favorable secondary structures of SC3 RRE variants and single mutants were insufficient to explain the functional activity differences observed between the V08-1and V20-1 RREs, we next assessed the ability of the RREs to adopt alternative conformers by evaluating their SHAPE-guided Shannon entropies and base pairing probability at single nucleotide resolution. Shannon entropies were calculated based on the probability for each base-pair appearing across all possible structures predicted for the RNA. Regions with highly stable well-defined RNA structures are characterized by lower Shannon entropies. Conversely, regions with high Shannon entropy are likely to form alternative conformers.

We compared Shannon entropy profiles of the V08-1, V09-2, V10-2, V20-1, M1, M2, M3, and M4 SC3 RREs (Figure 8). Most regions of these RREs exhibited low Shannon entropy, suggesting their overall secondary structures are highly stable. This result is consistent with the observation that these RRE RNAs migrate as a single discrete band on a native agarose gel, suggesting a high degree of structural homogeneity (data available upon request).

**Figure 8:**
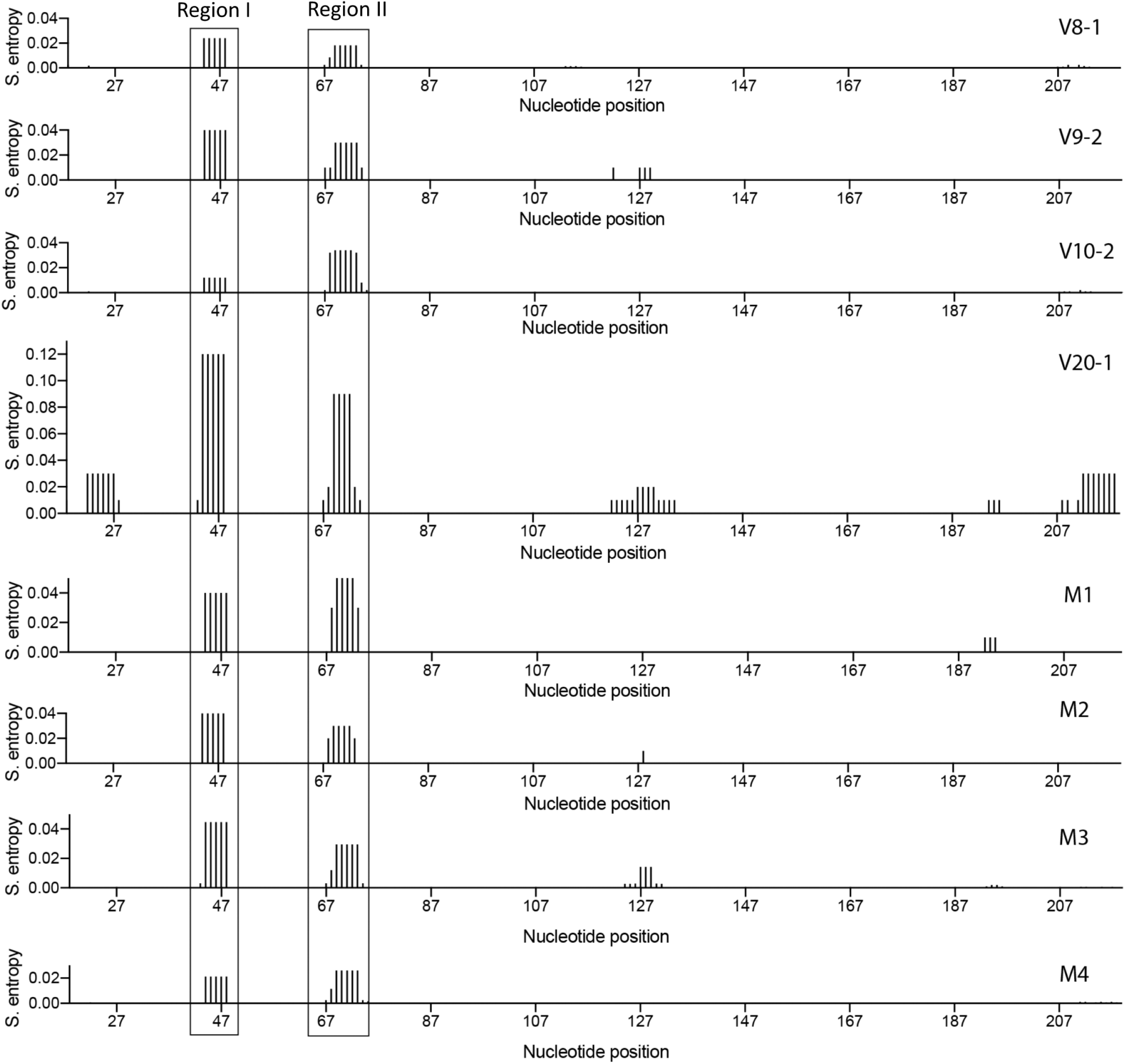
Shannon entropy profiles of SC3 RRE variants and single mutants. Shannon entropy values of the SHAPE-MaP generated RRE structures, smoothed over centered 11-nt sliding windows are plotted as a function of nucleotide position. Higher Shannon entropy suggests regions that are structurally dynamic. The boxed regions correspond to the loop between SL-IIA and SL-IIB (Region I) and the loop between SL-IIB and SL-IIC (Region II). These regions display high Shannon entropy in the high-activity V20-1 structure and low entropy in the lower activity V08-1 structure.

However, we observed differences in Shannon entropy values at two different regions. Specifically, values of the loop regions between SL-IIA and SL-IIB (Region I) and between SL-IIB and SL-IIC (Region II) varied between RREs. The values for the V20-1 RRE were, respectively, 5-6 fold and 4 fold higher than for the V08-1 and V10-2 RREs, suggesting that the nucleotides in these two regions of the V20-1 RRE are not always single-stranded as depicted in the secondary structures generated by SHAPE-MaP. This finding is further corroborated by SHAPE-MaP-guided base pairing probability calculated for each of these RREs (data available upon request), which shows that the two higher entropy regions of V20-1 RRE can base-pair with each other with a probability of 10-80%. Both the Shannon entropy and pairing probability data suggest that these two regions of V20-1 RRE are structurally dynamic, a feature that confers accessibility to protein interaction with the RNA by serving as landing pads for protein cofactors (44). This accessibility may have implications for Rev-binding, possibly contributing to the higher activity of V20-1 RRE.

Furthermore, both the V09-2 and M3 RREs have 2-fold higher Shannon entropy for the loop region between SL-IIA and SL-IIB (Region I) compared to V08-1 and V10-2 RREs. Their entropy values for the nucleotides between SL-IIB and SL-IIC (Region II), however, did not change relative to V08-1 and V10-2. While both V09-2 and M3 undergo the structural shift characteristic of V20-1 and are modestly more active than V08-1 and V10-2, the inability of V09-1 and M3 RREs to reach the degree of structural flexibility of the V20-1 RRE in the internal loop of SL-II might explain why these RREs are not as functionally active as V20-1. The Shannon entropy profile of M2 RRE is similar to the M3 and V09-1 RREs, whereas M1 RRE has 2-fold higher Shannon entropy than V08-1 and V10-2 in both regions of SL-II. The modestly higher entropy in the M1, M2, and M3 RREs relative to V08-1 and V10-2 might help to explain their trend towards higher functional activity (Figure 6). The M4 RRE, on the other hand, did not show any increase in entropy over V08-1 and V10-2 and also displayed similar or lower functional activity.

Taken together, our analysis suggests that both the major structural shift observed between the V08-1 and V20-1 RREs, and possibly also that the difference in structural dynamism in the internal loop of SL-II contribute to the significant increase in functional activity over time.

## DISCUSSION

In this study, we describe a complex relationship between RRE structure and functional activity using naturally occurring sequences obtained from a single patient. This study is the first detailed examination of longitudinal evolution of the HIV-1 RRE over about six years of naturally occurring infection and demonstrates clear selection pressures acting on the RRE sequence with a tendency towards increased functional activity over time. Increased activity can be explained by large-scale conformational changes within the RRE and a decrease in base-pairing stability at the initial Rev-binding site in SL-II.

The evolution of the RRE sequence is strongly suggestive of selection pressures operating on the RRE itself. The single founder RRE sequence found at visit 8 diversifies initially, and by visit 20 converges on only a single circulating RRE sequence differing from V08-1 by five single nucleotides. The early appearance and persistence of mutations at only a few locations, as well as the reduced viral diversity by visit 20, argue that the phenotype resulting from these changes was subject to strong selection pressures. Functional activity analysis of the SC3 RREs suggests that the RREs with higher activity were selected as disease progressed, with the early time-point RREs (V08-1 and V10-2) displaying the lowest and the late time-point RRE (V20-1) displaying the highest functional activity. The tendency towards increased functional activity corresponded with gradual accumulation, in different combinations, of five mutations that characterize V20-1 relative to V08-1.

There is no direct correspondence between specific mutations and functional activity phenotypes. Changes at the mutation 3 position predicted to disrupt base-pairing appear to result in an activity increase. However, V09-2 RRE containing this change alone still has significantly lower activity than V20-1, showing that the other mutations also contribute to a change in RRE activity in combination. While all mutations 1-5 are necessary to achieve the highest activity, their contributions are not merely additive. For example, V11-1 resembles V09-2 with the addition of mut 2, but it does not have increased activity relative to the founder. Additionally, mutations which have no impact on activity in isolation can increase activity in combination. For example, mut 4 does not change activity alone, but is responsible for the increase in activity between V14-2 and V20-1 in the context of the other four mutations.

Two types of structural changes were uncovered in this study that can explain differences in activity: major conformational changes and regions of increased base-pairing entropy. Previously, we have shown that alternative conformations of the laboratory strain NL4-3 RRE display significantly different activities, with a four stem-loop conformation supporting lower replicative capacity than a five stem-loop structure (21). In this study, we found that the higher activity RRE (V20-1) forms a five stem-loop conformation resembling that observed for NL4-3, while the lower activity RREs (V08-1 and V10-2) adopt a different conformation with a more collapsed central loop. As expected from functional analysis of the M3 and V09-2 RREs, both the G to A (V09-2) and G to C (M3) changes at the mutation 3 position induce a five stem-loop structure associated with increased activity. The additional mutations were not individually associated with large-scale conformational changes in SHAPE-MaP modeling.

The combination of mutations 1-5, though not each mutation in isolation, also corresponds to increased Shannon entropy in stem-loop II. This is the region where the first and second dimers of Rev bind (56). The disordered base pairing in this region, implied by greater entropy, may facilitate Rev-RNA binding at the primary Rev binding site and subsequent multimerization leading to enhanced activity. This is consistent with our previous *in vitro* gel-shift observations (43) that demonstrated the ability of the V20-1 RRE to promote Rev multimerization more efficiently than the V10-2 RRE. In the absence of major conformational changes attributable to mutations 1, 2, 4, and 5, this mechanism may account for the difference in functional activity between the RREs containing unpaired nucleotides at mutation 3 position (e.g. V09-2 and M3) that lack these additional mutations and the V20-1 RRE. It remains unclear how the presence of these mutations in combination results in increased stem-loop II entropy.

This study also provides further support for the hypothesis that the Rev-RRE regulatory axis plays an important role in HIV pathogenesis. An important consideration in the interpretation of these findings is that all five mutations in V20-1 relative to V08-1 are nonsynonymous in the overlapping gp41 coding sequence, raising the possibility that changes in Env could contribute to the selective pressure. We searched for known CTL epitopes in the Env ORF overlapping the V08-1 RRE sequence using the Los Alamos HIV database Epitope Location Finder tool (www.hiv.lanl.gov). CTL epitopes have been described in the V08-1 regions containing three of the five mutations (mutations 3, 4, and 5). Thus, selection of these mutations could plausibly reflect immune evasion, though it is notable that neither mutation 3 nor 5 were consistently found in all RRE sequences at all time points after they were first observed at visits 9 and 10.

While the selection evidenced by sequence convergence at visit 20 might be due to pressure on Env in terms of immune evasion, replication efficiency, or other factors, it is likely that functional differences in Rev-RRE activity also play a significant role in overall viral fitness. Nucleotide changes at the mutation 3 position highlight this likelihood. The nonsynonymous mutations from G to A at visit 9 and A to C at visit 19 are both consistent with CTL evasion. From the standpoint of the RRE, however, the G to A change shifts the RRE structure to a higher activity conformation while A to C is structurally and functionally neutral. Thus, the A to C mutation preserves high RRE activity while also modifying the Env amino acid sequence. Convergence of circulating RRE sequences by visit 20 is particularly striking in light of the very low CD4 count at this time point, as viral fitness would likely have been more influenced by Rev-RRE activity than by immune surveillance.

We were unable to assess functional evolution of the Rev protein over time in this study due to the sequencing strategy used, which created reads too short to allow linkage of both of the Rev coding exons and the RRE. However, previous work using single genome sequencing with samples from this patient (43) demonstrated that a single Rev amino acid sequence was most prevalent at both visits V10 and V20. This is the Rev sequence that was used in this study. Its persistence throughout the entire infection period suggests that it likely occurred in conjunction with most or all of the RREs tested here.

It is also notable that m6A methylation of RRE RNA has recently been proposed as an additional mechanism of modulating HIV replication capacity (57). This was not assessed in the present study but could additionally contribute to differences in Rev-RRE activity.

The generalizability of the findings described here awaits further longitudinal studies, analyzing both Rev and RRE sequences, in patients with varying courses of disease. A clear understanding of RRE evolution during natural infection will help to further our understanding of the role that this regulatory axis may play in key aspects of the viral life cycle, including transmission and the establishment of latency. Furthermore, since the Rev-RRE interaction is an essential step in viral replication, it is also a promising target for drug development. This study demonstrates that RRE structures can vary over the course of natural infection, and thus that rational drug design must account for the spectrum of potential Rev-RRE interaction conformations.

## MATERIALS AND METHODS

### Clinical samples and ethics statement

All studies involving human subjects were approved by the institutional review board at Montefiore Hospital, Bronx, NY, as part of the Women’s Interagency HIV Study (WIHS). Written informed consent was provided by all study subjects. Plasma samples from one patient (SC3), collected over a period of seven years, were obtained from the WIHS Consortium, Bronx, NY (Kathryn Anastos, principal investigator), as previously described (43). A total of 11 samples representing different time points were received for analysis.

### Viral RNA extraction and cDNA synthesis

Viral RNA was extracted from 1 ml of plasma using a guanidinium extraction protocol (58). Viral cDNA synthesis was performed immediately after viral RNA was extracted as described (43). Briefly, each viral RNA pellet was resuspended in 40 µl of 5 mM Tris-HCL, pH 8.0, prior to cDNA synthesis with 50µM of oligo(dT) and 100 U of Superscript III per reaction (Invitrogen, Carlsbad, CA). The reverse transcription reaction was incubated at 50°C for 1 hour, then 55°C for 1 hour and then 70°C for 15 minutes, after which 4 units of RNase H (Invitrogen) were added, and samples incubated at 37°C for 20 minutes. Samples were stored at −80°C prior to amplicon generation.

### PCR amplification and purification

Amplicons (~3 kb) spanning both HIV Rev exons and the RRE, were produced for each time point. The first round PCR reaction was performed by adding 1 µl of cDNA template to a 20 µl reaction containing 0.005 U of Platinum Taq Hi-Fidelity polymerase (Invitrogen) as previously described (59). Using 1 µl of the first round PCR product as a template, nested PCR resulted in amplicons ~3 kb in length. Primers for the first round PCR reaction were 2302 (5’-aagccacctttgcctagtg-3’) and 2278 (5’-ttgctacttgtgattgctccatgt-3’) while those for the nested PCR were 2277 (5’-tagagccctggaagcatccaggaag-3’) and 2280 (5’-gtctcgagatactgctcccaccc-3’) (43). Amplicon size was verified by agarose gel electrophoresis.

Each 20 µl PCR reaction was purified with 36 µl of AMPure XP beads (Agencourt). After addition of beads, samples were pipetted up and down 10 times and incubated at room temperature for 10 minutes prior to being placed on a magnetic stand. After a 2 minute incubation, supernatant was removed and the beads were washed twice with 80% ethanol. After beads were air-dried on a magnetic stand for 10 minutes, they were removed from the magnet, resuspended in 52.5 µl of elution buffer, and incubated at room temperature for 2 minutes. Beads were then placed on the magnetic stand for 2 minutes, or until the supernatant was cleared. The cleared supernatant containing the purified PCR amplicons was removed and placed in a new 96-well plate then stored at −20°C.

### DNA library preparation and sequencing

Prior to preparing the DNA library, purified amplicon products were quantified with the dsDNA HS (high sensitivity) assay kit (Invitrogen) using the Qubit 2.0 fluorometer (Invitrogen). The DNA library was prepared using the standard Nextera XT DNA library preparation protocol (Illumina). A total of 1 ng of input DNA for each sample was added to the reaction buffer which fragments the input DNA and adds adapter sequences to the ends to allow for PCR reactions downstream in the library preparation process. A brief PCR cycle followed the fragmentation protocol to add indexes used to identify each individual sample and additional sequences for proper cluster formation. PCR samples were purified and size-selected for 300-500 bp amplicons using 25 µl of AMPure XP beads. Amplicons were pooled together in equimolar concentrations then sequenced with the Illumina MiSeq Reagent Kit v3 (600 cycles).

### RRE sequence analysis

Paired end reads for each time point were generated using an Illumina MiSeq. The two overlapping paired reads were merged using FLASH (60), a plugin in Geneious (Biomatters, Auckland, New Zealand) to produce a single read for each pair. These reads were then aligned to the NL4-3 genome and reads overlapping the RRE were extracted. Extracted reads were filtered for individual reads that spanned the full length of the short RRE. These reads were assembled into contigs using the Geneious *de novo* assembly tool, set for a minimum overlap identity of 100% and a minimum overlap of the entire 234-nt minimal RRE. The frequency of each contig was calculated based on the number of reads assigned to each contig. Contigs comprising less than 5% of the total reads were then removed from each time point (to eliminate minor HIV species and/or potential PCR errors) and the frequencies of the remaining contigs were recalculated based on the total number of reads remaining. The sequence of each contig was then aligned to the sequence from the major V08 contig to generate the alignment shown in Figure 2 (see also Figures S1 and S2). RREs were named using the format VXX-Y where XX corresponds to the WIHS cohort visit number at which the plasma sample generating the sequence was obtained and Y corresponds to the frequency rank order of the sequence. Thus, V10-2 refers to the second most prevalent RRE from the sequencing of the plasma sample obtained at WIHS visit 10.

### *In vitro* transcription of RRE RNAs

RRE RNAs were *in vitro* transcribed by T7 RNA polymerase using the MEGAshort-script kit (Life Technologies) per manufacturer’s guidelines. Templates for *in vitro* transcription were generated by PCR amplification of RREs from plasmids containing synthetic sequences (Integrated DNA technologies) using oligonucleotides designed to introduce both a T7 promoter sequence at the 5’ end of the 234-nt and 351-nt RREs and a structure cassette at the 3’ end of the 351-nt RREs (Table S1). Plasmids used for generating transcription DNA template for the 234-nt V08-1, V09-2, V10-2, V20-1, M1, M2, M3, and M4 RREs were designated pHR 5334, pHR 5766, pHR 4784, pHR 4788, pHR 5326, pHR 5328, pHR 5330, and pHR 5332, respectively. Those used to generate 351-nt V10-2 and V20-1 RREs were designated pHR 5321 and pHR 5322, respectively. *In vitro* transcription reactions were treated with Turbo DNase I (Life Technologies) for 1 hr at 37°C to digest the DNA template, heated to 85°C for 2 min, and RNA was fractionated on a denaturing gel (5% polyacrylamide (19:1), 1x TBE, 7M urea) at constant temperature (45°C, 30W max). RRE bands were located by UV shadowing, excised, electroeluted at 200 V for 2 hours at 4°C, ethanol precipitated, and stored at 4°C in TE buffer (10 mM Tris, pH 7.6; 0.1 mM EDTA).

### Folding and SHAPE-MaP profiling of RRE RNA

For each RNA, 1M7 (+), 1M7 (−), and denaturation control reactions were generated. Approximately 5 pmoles of RRE RNA was incubated with renaturation buffer (1X: 10 mM Tris, pH 8.0, 100 mM KCl, 0.1 mM EDTA) in a total volume of 5 μl, heated to 85°C, and renatured by slow cooling (0.1°C/sec) to 25°C. Renatured RNA was incubated with RNA folding buffer (1X: 35 mM Tris pH 8.0, 90 mM KCl, 0.15 mM EDTA, 5 mM MgCl2, 2.5% glycerol) in a total volume of 9 μl at 37°C for 30 min. RNA was modified by addition of 1 µL of 25 mM 1M7. The negative control reaction contained 1 µL of DMSO instead of 1M7. The denaturation control was produced by incubating 5 pmoles of RNA with denaturation buffer (1X: 50% formamide, 50 mM HEPES, and 4 mM EDTA) in a volume of 9 µL at 95 °C for 1 min. 1 µL of 20 mM 1M7 was added and the mixture incubated at 95 °C for another 1 min. RNA from 1M7 (+), 1M7 (−), and denatured control reactions were recovered by ethanol precipitation, resuspended in TE buffer (10 mM Tris, pH 7.6; 0.1 mM EDTA), and stored at −20°C.

### Mutational profiling of modified RNA

RNA from 1M7 (+), 1M7 (−), and denatured control reactions were reverse transcribed using corresponding oligos to generate a cDNA library (Table S1). Reverse transcription was performed by first annealing 2 µM of RT oligo to the RNA in a reaction volume of 11 µL by incubation at 65 °C for 5 min, followed by cooling on ice. cDNA synthesis was initiated by incubating the annealing mixture with 8 µL of 2.5X RT reaction mixture (2.5X: 125 mM Tris (pH 8.0), 187.5 mM KCl, 15 mM MnCl_2_, 25 mM DTT and 1.25 mM dNTPs) and 1 µL of Superscript II RT (Thermo Fisher, 200 U/µL) for 42 °C for 3 hours. The RNA template was then hydrolyzed by adding 1 µL 2 N NaOH, followed by neutralization by addition of 1 µL 2N HCl. The cDNA library was purified using G50 spin columns (GE Healthcare).

For mutational profiling, cDNA was converted to dsDNA with Illumina adapters for high throughput sequencing on an Illumina platform. This was achieved in two consecutive PCR reactions, namely PCR1 and PCR2. The first reaction (PCR1) appended partial Illumina adapters to the ends of the amplicons. The entire cDNA library was used as template in a 100 µl PCR1 reaction (1.1 µL each of 50 pmoles of forward and reverse oligo, 2 µL of 10 mM dNTPs, 20 µL of 5x Q5 reaction buffer, 1 µL of Hot Start High-Fidelity DNA polymerase (NEB)) using cycling conditions: 98°C for 30 sec, 15 cycles of (98 °C for 10 sec, 50 °C for 30 sec, 72 °C for 30 sec), 72 °C for 2 min. PCR product was gel purified and 10% was used as template DNA in the PCR2 reaction. The PCR2 completed the Illumina adapters while adding appropriate barcoding indices. The PCR2 reaction and cycling conditions were as those for PCR1. The resulting amplicon library was fractionated through a 2% agarose gel and amplicons purified from the gel slices by electro-elution at room temperature for 2 hr followed by ethanol precipitation. Each amplicon library was quantified by real time PCR using the KAPA Universal Library Quantification Kit (Illumina) per manufacturer’s protocol. Sequencing libraries were pooled and mixed with 20% PhiX and sequenced on an Illumina MiSeq to generate 2 × 150 paired-end reads. Sequence files were fed into ShapeMapper (v1.2) software (49) to generate SHAPE reactivity profiles by aligning the reads to 234-nt RRE RNA sequence using the software with default settings. Reactivity values obtained from ShapeMapper were input to RNAstructure software (51) to generate the minimum free-energy RNA secondary structure model.

### RRE gel migration assay

RRE structural homogeneity was assessed by comparing the migration rate of the folded RREs on native agarose and polyacrylamide gels. Approximately 20 pmoles of RNA was suspended in 5 μl renaturation buffer (1X: 10mM Tris, pH 8.0, 100mM KCl, 0.1mM EDTA), heated to 85°C, and renatured by slow cooling (0.1°C/sec) to 25°C. Renatured RNA was incubated with 5 μl of 2X RNA folding buffer (1X:70 mM Tris pH 8.0, 180 mM KCl, 0.3 mM EDTA, 10 mM MgCl_2_, 5% of glycerol) at 37°C for 30 min, and fractionated through a native 8% polyacrylamide gel (29:1) run for 22 hours at 4°C or on a 2% native agarose. RRE bands were visualized by UV shadowing and ethidium bromide staining, respectively.

### Plasmid constructs for functional assays of Rev-RRE activity

Functional activity of Rev-RRE pairs was determined by means of a fluorescence-based transient transfection assay. To test RRE sequences, a plasmid containing a near full length NL4-3 HIV-1 sequence (61) was modified to express blue fluorescent protein (BFP) in a Rev-independent fashion and green fluorescent protein (GFP) in a Rev-dependent fashion. The native 351-nt RRE within *env* was flanked by XmaI and XbaI sites to permit exchange of this sequence with other RREs of interest. Additional modifications were made to ensure the construct was not replication-competent and could not express Rev. To create additional RRE constructs, the native RRE was removed by digesting the plasmid with XmaI and XbaI. Commercially synthesized 234-nt RRE sequences of interest with appropriate flanking sequences (Integrated DNA Technologies) were then cloned into the opened *env* region using Gibson assembly.

Rev was provided *in* trans using one of two constructs. The predominant Rev previously identified at V10 and V20 by single genome sequencing had identical amino acid sequences. This Rev was designated M0-B/M57-A and used for functional assays except as otherwise noted (43).

For the analysis described in Figure 4, the M0-B/M57-A *rev* sequence was cloned into a CMV expression plasmid. For the remaining functional assays, a separate Rev-expressing construct was created by modifying the murine stem cell virus construct, pMSCV-IRES-mCherry FP (Addgene plasmid # 52114), to express both the M-0B/M57-A Rev and the mCherry fluorescent protein by means of a bicistronic transcript including an internal ribosome entry site (62). The MSCV vector was a gift from Dario Vignali (unpublished). Constructs utilized in the functional assays are listed in Table S2.

### Selection of RREs for functional analysis

Both naturally occurring RREs sequenced from patient SC3 and synthetic RREs containing individual mutations of interest were tested for functional activity. All RREs tested were 234-nt in length. The naturally occurring RREs assessed were V08-1, V09-2, V10-2, V11-1, V13-1, V14-2, V19-2, and V20-1. Additionally, synthetic RREs were created based on single nucleotide differences between the sequences of V10-2 and V20-1. The mutation 1 RRE (M1) is identical to V10-2 with the exception of a G to A change at position 61, reflecting the G to A change seen at that position in V20-1. The mutation 2 RRE (M2) is identical to V10-2 with the exception of a T to A change at position 84. Of note, this RRE sequence was also observed to occur naturally as sequence V12-1. The mutation 3 RRE (M3) is identical to V10-2 with the exception of a G to C change at position 194. The mutation 4 RRE (M4) is identical to V10-2 with the exception of an A to G change at position 228.

### Rev-RRE functional activity assays

RRE-containing constructs were designed such that GFP expression occurs in a Rev-dependent fashion while BFP expression occurs in a Rev-independent fashion. In cells transfected with both the RRE-containing and Rev-containing constructs, the degree of GFP expression relative to BFP expression is proportional to the functional activity of the tested Rev-RRE pair (Jackson et al.. manuscript in preparation).

Except as noted, transfections were performed using the polyethylenamine method and 8×10^5^ 293T/17 cells in each well of a 12-well plate. Cells were maintained in 1 mL IMDM supplemented with 5% bovine calf serum. Each RRE-containing construct was tested individually with the SC3 Rev-containing construct. Additionally, each RRE-containing construct was transfected into cells without the addition of Rev to ensure that GFP expression in this system was truly Rev-dependent. In transfections performed without the Rev-containing construct, an empty CMV construct was used to compensate such that each transfection was performed with a constant mass of plasmid. For each transfection, 1000 ng of an RRE-containing construct and 100 ng of either the MSCV-Rev construct or empty vector was used. Simultaneously, different cultures of 293T/17 cells were transfected with constructs expressing GFP, BFP, or mCherry alone as single-color controls to permit color compensation during flow cytometry. Each set of transfections was performed in duplicate with the exception of V09-2 and V13-1, for which four replicates were performed. After transfection, cells were incubated for 48 h, then suspended in phosphate buffered saline. Flow cytometry was performed on the resulting suspension using an Attune NxT flow cytometer with autosampler attachment (Thermo Fischer Scientific). Data acquisition was performed on the Attune NxT software package using the following channels:

|  |  |  |  |
| --- | --- | --- | --- |
| Laser line | 488 | 405 | 561 |
| Emission filter | 530/30 | 440/50 | 620/15 |
| Fluorescent protein | eGFP | TagBFP | mCherry |

Post-acquisition color compensation and data analysis was performed using FlowJo v10 (FlowJo, LLC). For each analysis, gating was performed on single 293T/17 cells. Next, a daughter population of cells that expressed the RRE-containing transcript was identified by gating on positivity for GFP or BFP. Using this population, the ratio of arithmetic mean fluorescent intensity (MFI) of eGFP to TagBFP for successfully transfected cells was calculated in both the presence and the absence of Rev. A low ratio in the absence of Rev and a high ratio in its presence indicated that GFP expression was Rev-dependent.

After determining that the RRE-containing constructs expressed GFP in a Rev-dependent fashion, an additional analysis was performed to determine the functional activity of specific Rev-RRE pairs. In this further analysis, only those transfections including both an RRE-containing construct and the Rev-containing construct were considered. An additional daughter population was created from that described above by gating on mCherry positivity. This final population consisted only of single 293T/17 cells expressing GFP and/or BFP and also mCherry. This pattern of expression ensured that selected cells were successfully co-transfected with both plasmid constructs. Once this population was defined, the ratio of GFP to BFP MFI was re-calculated. This final GFP:BFP ratio was used to determine the relative functional activity of different RREs assessed in the presence of SC3 Rev. Differences in functional activity values between RREs were evaluated using SPSS Statistics v25 (IBM). P-values were calculated using a two-tailed one-way ANOVA with Tukey’s HSD test for Figures 4 and 7 and with a one-tailed one-way ANOVA with Dunnett’s test for Figure 6E.

Analysis of the V08-1, V10-2, and V20-1 RREs described in Figure 4 was performed with minor modifications. For this analysis, Rev was provided *in trans* by the CMV-SC3 Rev construct. Each well of a 12-well plate was seeded with 4×10^5^ 293T/17 cells maintained in 2 mL IMDM supplemented with 10% bovine calf serum. One day after plating, medium was removed from each well and replaced with 1 mL IMDM supplemented with 5% bovine calf serum. Cells in each well were then transfected with 1000 ng of an RRE-containing construct and 50 ng of the CMV-Rev construct using the polyethylenamine method. Four transfection replicates were performed. Simultaneously, different cultures of 293T/17 cells were transfected with constructs expressing GFP or BFP alone as single-color controls to permit color compensation during flow cytometry. Cells were harvested and flow cytometry was performed 24 hours after transfection. Gating was performed as above identifying cells successfully transduced with the RRE-containing construct and expressing GFP or BFP, and the ratio of GFP:BFP was calculated. Additional gating on mCherry positive cells was not performed.

### CE-SHAPE

RNA was folded as in the gel migration assay except that the final volume of the folded RNA mix was 150 ul and glycerol was excluded from the folding buffer. Folded RNAs were probed using 3 mM NMIA. For this, RNAs were divided into experimental (NMIA+) and control (NMIA−) aliquots (72 μl each), to which 8 μl 30 mM NMIA in anhydrous DMSO or DMSO alone was added, respectively. Modification reactions were incubated at 37°C for 50 min, ethanol precipitated, and re-suspended in 13 ul nuclease-free water. Reverse transcription of modified RNAs, cDNA processing/fractionation, and SHAPE data analysis were conducted as previously described (63).

### Creating probability pairing and Shannon entropy profiles

Pairing probability (Figure S5-S8) and Shannon entropy values were calculated by feeding SuperFold (49) with the 1M7 reactivities of the RREs. Shannon entropy measurements were calculated over a centered 11-nt sliding window and plotted against nucleotide position.

### Nucleotide sequence accession numbers

Nucleotide sequences generated through the procedure above corresponding to RREs from viral quasispecies found in samples from patient SC3 were deposited in Genbank under accession numbers MK190736 through MK190867. The SC3 Rev sequence used in the functional assays was previously deposited under accession number KF559146. The RRE sequences listed here as V10-2 and V20-1 were previously deposited under accession numbers KF559160 and KF559162, respectively.

## ACKNOWLEDGMENTS

This work was supported by grants GM 110009, AI134208, AI087505 and AI068501 as well as the Women’s Interagency HIV Study (WIHS) grant UO1-AI-35004 from the National Institutes of Health (NIH). The patient samples used in this manuscript were collected by the Women’s Interagency HIV Study (WIHS). The Bronx WHIS repository provided the initial patient materials and the WIHS central repository provided the materials from the intermediate time points (Bronx WIHS [Kathryn Anastos and Anjali Sharma], U01-AI-035004). Jason W Rausch (NCI) is acknowledged for his contribution in setting up the SHAPE-MaP analysis pipeline. S.L.G. and C.S. were supported by the Intramural Research Program of the National Cancer Institute, National Institutes of Health, Department of Health and Human Services. P. E. H. J. was supported by grant K08AI136671 from the National Institutes of Health. Salary support for M.-L.H. and D.R. was provided by the Charles H. Ross, Jr., and Myles H. Thaler Endowments at the University of Virginia. The contents of this publication are solely the responsibility of the authors and do not represent the official views of NIH or the WIHS. The funders had no role in study design, data collection and interpretation, or the decision to submit the work for publication.

