## Supplemental Figures for "Evolution of the HIV-1 RRE during natural infection reveals key nucleotide changes that correlate with altered structure and increased activity over time"

Figure S1: Comprehensive evolutionary alignment of SC3 RREs. The prevalence of each RRE sequence at a given visit was calculated as a percentage of the total number of sequences generated from that plasma sample without removing minority variants. RRE sequences were assigned a code in the form VXX-Y where XX refers to the visit number and Y refers to the prevalence rank order of that sequence. Nucleotide differences relative to the V08-1 presumptive founder sequence are highlighted. Sequences in red boxes have a prevalence of at least 5% and were included in the analysis presented in Figure 2 after excluding the minority variants.

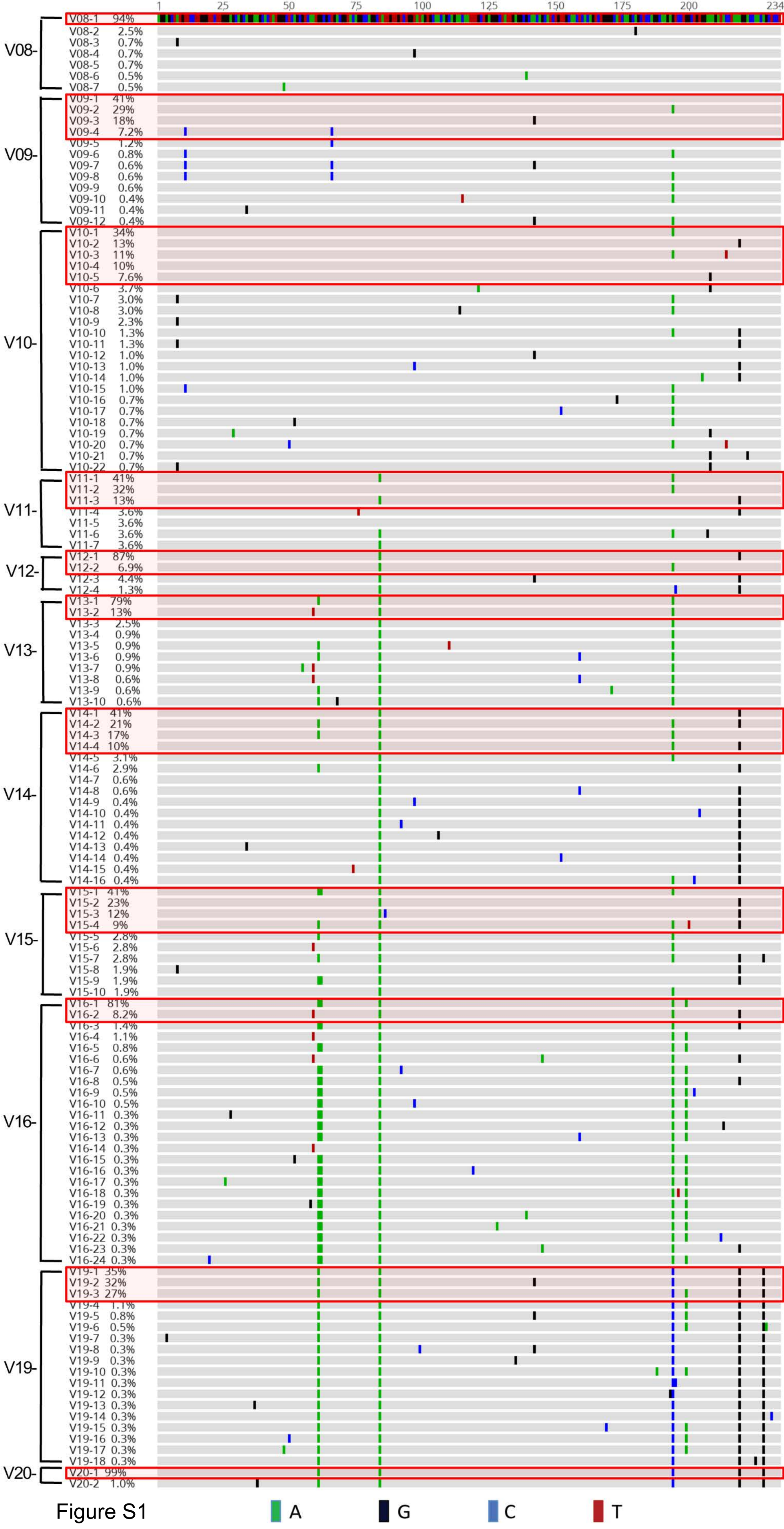
1
25
50
75
100
125
150
175
200
234
V08-
V08-1 94%
V08-2 2.5%
V08-3 0.7%
V08-4 0.7%
V08-5 0.7%
V08-6 0.5%
V08-7 0.5%
V09-
V09-1 41%
V09-2 29%
V09-3 18%
V09-4 7.2%
V09-5 1.2%
V09-6 0.8%
V09-7 0.6%
V09-8 0.6%
V09-9 0.6%
V09-10 0.4%
V09-11 0.4%
V09-12 0.4%
V10-
V10-1 34%
V10-2 13%
V10-3 11%
V10-4 10%
V10-5 7.6%
V10-6 3.7%
V10-7 3.0%
V10-8 3.0%
V10-9 2.3%
V10-10 1.3%
V10-11 1.3%
V10-12 1.0%
V10-13 1.0%
V10-14 1.0%
V10-15 1.0%
V10-16 0.7%
V10-17 0.7%
V10-18 0.7%
V10-19 0.7%
V10-20 0.7%
V10-21 0.7%
V10-22 0.7%
V11-
V11-1 41%
V11-2 32%
V11-3 13%
V11-4 3.6%
V11-5 3.6%
V11-6 3.6%
V11-7 3.6%
V12-
V12-1 87%
V12-2 6.9%
V12-3 4.4%
V12-4 1.3%
V13-
V13-1 79%
V13-2 13%
V13-3 2.5%
V13-4 0.9%
V13-5 0.9%
V13-6 0.9%
V13-7 0.9%
V13-8 0.6%
V13-9 0.6%
V13-10 0.6%
V14-
V14-1 41%
V14-2 21%
V14-3 17%
V14-4 10%
V14-5 3.1%
V14-6 2.9%
V14-7 0.6%
V14-8 0.6%
V14-9 0.4%
V14-10 0.4%
V14-11 0.4%
V14-12 0.4%
V14-13 0.4%
V14-14 0.4%
V14-15 0.4%
V14-16 0.4%
V15-
V15-1 41%
V15-2 23%
V15-3 12%
V15-4 9%
V15-5 2.8%
V15-6 2.8%
V15-7 2.8%
V15-8 1.9%
V15-9 1.9%
V15-10 1.9%
V16-
V16-1 81%
V16-2 8.2%
V16-3 1.4%
V16-4 1.1%
V16-5 0.8%
V16-6 0.6%
V16-7 0.6%
V16-8 0.5%
V16-9 0.5%
V16-10 0.5%
V16-11 0.3%
V16-12 0.3%
V16-13 0.3%
V16-14 0.3%
V16-15 0.3%
V16-16 0.3%
V16-17 0.3%
V16-18 0.3%
V16-19 0.3%
V16-20 0.3%
V16-21 0.3%
V16-22 0.3%
V16-23 0.3%
V16-24 0.3%
V19-
V19-1 35%
V19-2 32%
V19-3 27%
V19-4 1.1%
V19-5 0.8%
V19-6 0.5%
V19-7 0.3%
V19-8 0.3%
V19-9 0.3%
V19-10 0.3%
V19-11 0.3%
V19-12 0.3%
V19-13 0.3%
V19-14 0.3%
V19-15 0.3%
V19-16 0.3%
V19-17 0.3%
V19-18 0.3%
V20-
V20-1 99%
V20-2 1.0%
A
G
C
T

Figure S1

Figure S2: Evolutionary alignment of SC3 RREs at single-nucleotide resolution. The alignment includes only contigs that were present in at least 5% of the sequences at a given time point as explained in the Methods. Sequences obtained from different plasma samples corresponding to different time points are separated by horizontal red bars.

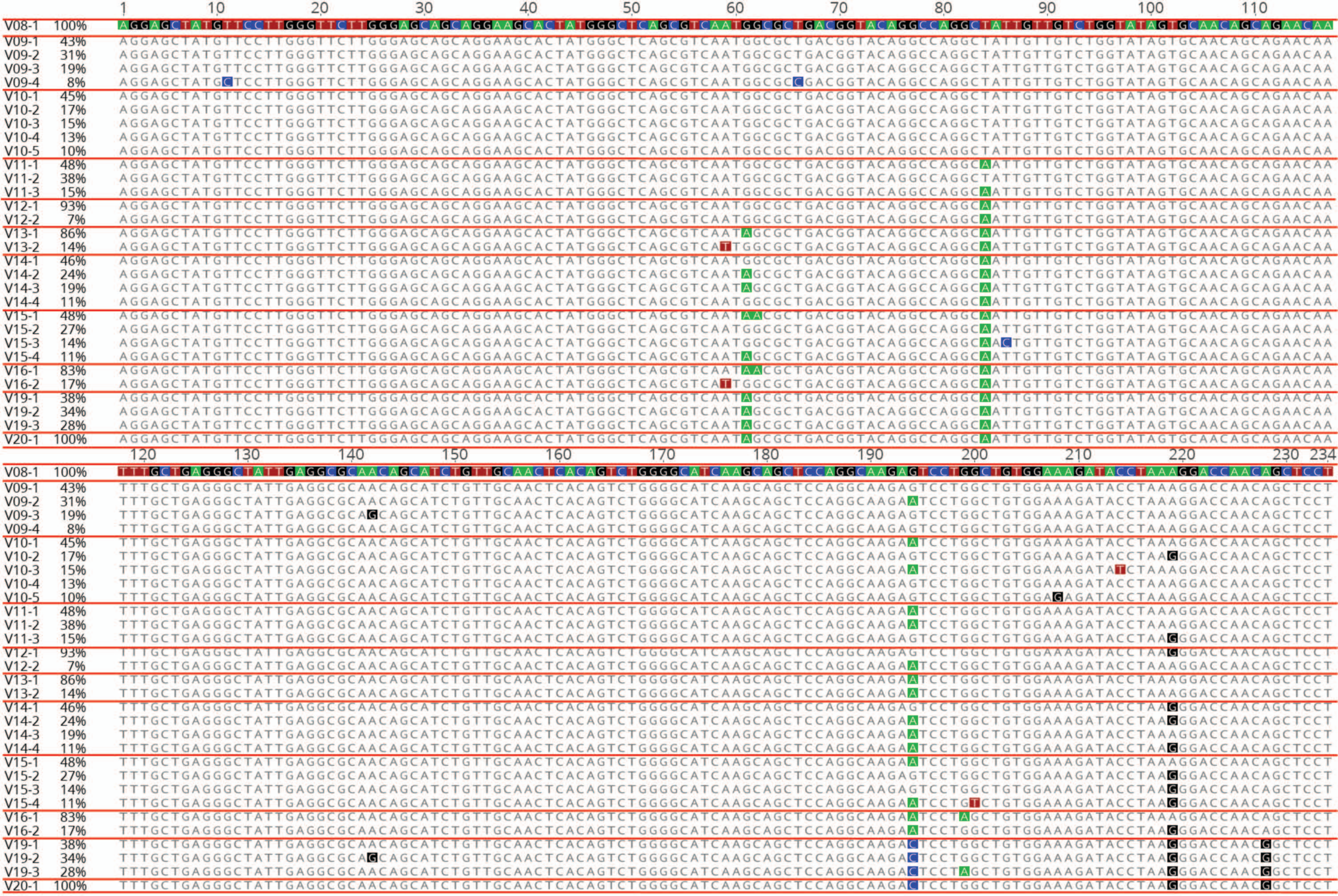
V08-1 100%
V09-1 43%
V09-2 31%
V09-3 19%
V09-4 8%
V10-1 45%
V10-2 17%
V10-3 15%
V10-4 13%
V10-5 10%
V11-1 48%
V11-2 38%
V11-3 15%
V12-1 93%
V12-2 7%
V13-1 86%
V13-2 14%
V14-1 46%
V14-2 24%
V14-3 19%
V14-4 11%
V15-1 48%
V15-2 27%
V15-3 14%
V15-4 11%
V16-1 83%
V16-2 17%
V19-1 38%
V19-2 34%
V19-3 28%
V20-1 100%
A
G
C
T

Figure S2
